# Machine Learning Reveals Key Glycoprotein Mutations and Rapidly Assigns Lassa Virus Lineages

**DOI:** 10.1101/2024.07.31.605963

**Authors:** Richard Olumide Daodu, Jens Uwe-Ulrich, Denise Kühnert

## Abstract

Lassa fever, caused by the Lassa virus (LASV), remains a major public health concern in West Africa, causing numerous fatalities annually and several intercontinental cases since its discovery in 1969. Despite ongoing research, no approved vaccines are available, with current efforts focusing on immunotherapy. LASV is divided into distinct lineages that circulate in specific geographic regions, elicit varying immune responses, and exhibit different pathophysiological effects. Understanding the genetic differences between these lineages is crucial for developing, improving, and distributing diagnostics, treatments, and vaccines. In this study, we analyzed the LASV glycoprotein complex (GPC), the only surface protein, using statistics, machine learning, and phylogenetics. At a population scale, we identified key amino acid differences between Nigerian lineages and those in other West African regions, particularly near the stable signal peptide cleavage site and other immune-related regions (e.g., AA positions 59–76). Additionally, we found that GPC sequences from Lineages II and III are shorter than those from Lineage IV, due to the high prevalence of a codon insertion at positions 178–180 (amino acid position 60). This insertion may contribute to inaccuracies observed in molecular diagnostics for LASV and may also play a role in the increased fatality associated with Lineage IV. The insertion has reemerged and persisted in Lineage II which may indicate a fitness advantage. Furthermore, we developed a fast and highly accurate lineage classification tool called CLASV that allows rapid identification of LASV lineages, improving the ability to monitor emerging outbreaks and exported cases.

## Introduction

Since its discovery in 1969, the viral hemorrhagic disease Lassa fever (LF) constitutes a major public health threat, with 100,000-300,000 cases and around 5,000 deaths annually in West Africa (AfricaCDC, n.d.). Although the overall fatality rate in hospitalized patients is relatively low (∼1%) (AfricaCDC, n.d.), it is very high among pregnant women (80%) and fetuses (up to 100%) (Agboeze et al., 2019; Salami et al., 2022). Currently, no approved vaccine nor appropriate therapeutic measures exist, although efforts are ongoing (Garry, 2023). The infectious agent causing Lassa fever is the Lassa Virus (LASV). Its main reservoir host is *Mastomys natalensis*, but several additional hosts with the potential to spread the virus having been discovered recently (Happi et al., 2023; Olayemi et al., 2016). Experimental research involving LASV is confined to Biosafety Level 4 (BSL4) laboratories, and the disease is currently on the World Health Organization’s (WHO) top priority list (WHO, n.d.). Exported LASV cases have been reported worldwide (Wolf et al., 2020), including in Europe and Asia, emphasizing the need for robust biosecurity measures, rapid diagnosis, effective treatments and vaccines.

To date, there are seven known LASV lineages circulating in distinct geographical locations (Figure 1A). Lineages I, II and III circulate in non-overlapping parts of Nigeria (Ehichioya et al., 2019), while lineage IV circulates in Sierra Leone, Guinea, and Liberia (Andersen et al., 2015), lineage V in Mali and Cote d’Ivoire (Manning et al., 2015), and lineage VII in Togo and Benin (Whitmer et al., 2018; Yadouleton et al., 2020). The LASV strain found in the host *Hylomyscus pamfi* in Nigeria was designated lineage VI (Olayemi et al., 2016; Whitmer et al., 2018).

**Figure 1:**
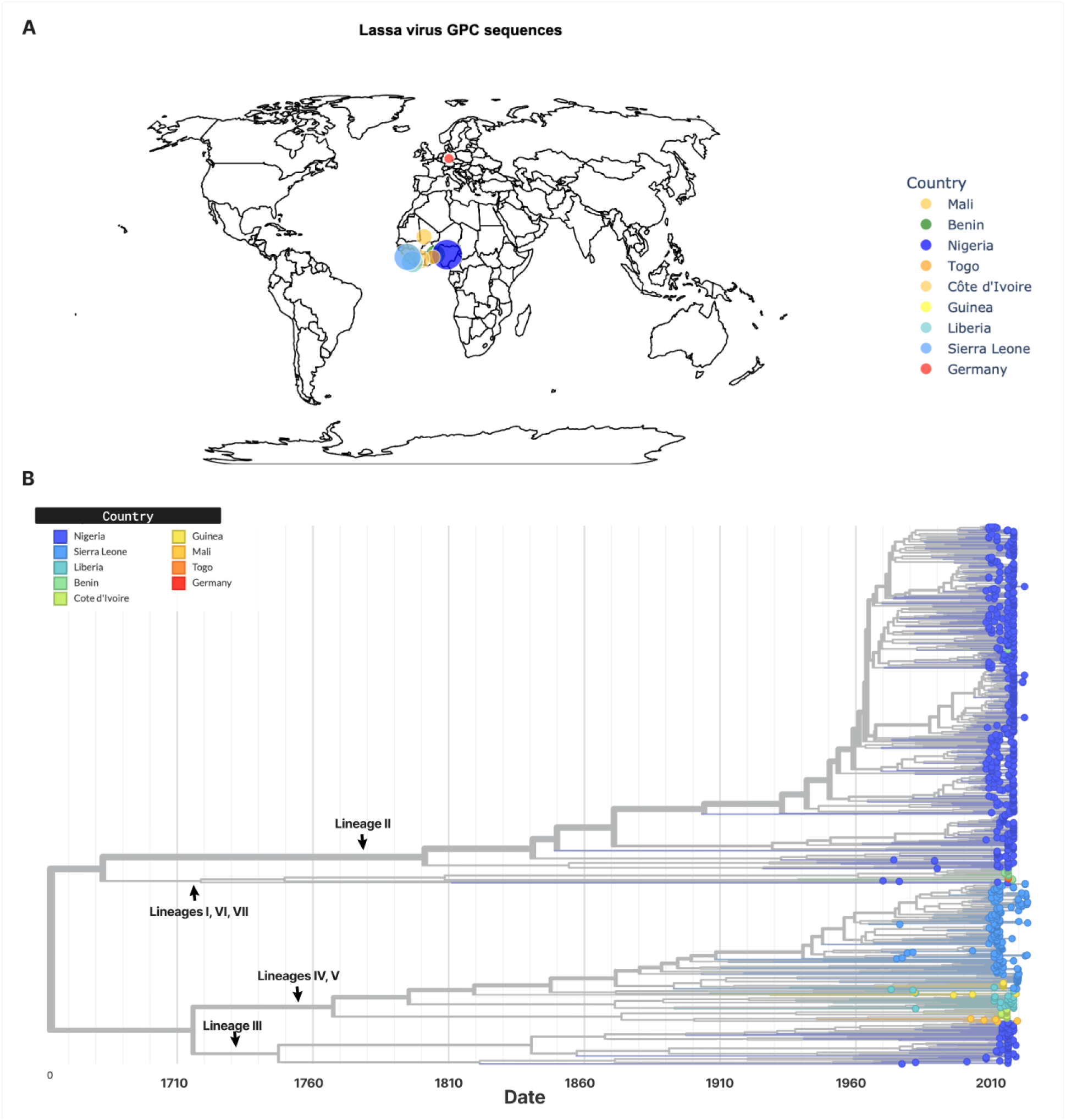
Lassa virus geographic distribution and phylogeny. (A) Geographic distribution of LASV glycoprotein sequences used in this study. Counts have been log-normalized. (B) Phylogenetic tree of the nucleotide sequences annotated by lineage (black arrows). Leaf nodes are coloured by geographical location.

Currently, treatment for Lassa fever relies primarily on ribavirin, a drug with severe side effects (De Franceschi et al., 2000; Salam et al., 2022). Immunotherapy is a promising treatment option. Early attempts at passive immunization included the use of convalescent plasma and immune serum, which showed some success (Clayton, 1977; Leifer et al., 1970). A monoclonal antibody cocktail, composed of three neutralizing antibodies targeting the LASV glycoprotein complex (GPC), has shown to be efficient in non-human primates against representatives of lineages I-IV (Mire et al., 2017). However, this antibody combination has been reported to confer heavily reduced protection against lineage VII (Woolsey et al., 2024). This may be due to mutations in the GPC, which may prevent binding of specific antibodies (Enriquez et al., 2022; Robinson et al., 2016). Thus, it is crucial to determine which lineages carry such mutations and to what extent. Unfortunately, comprehensive population scale genetics studies of LASV are scarce.

Significant differences in Lassa fever outcomes have been observed across various regions of West Africa. Specifically, LASV lineage IV, which circulates in Sierra Leone, has been reported to be considerably more lethal than the Nigerian lineages (Andersen et al., 2015; Akhuemokhan et al., 2017). Investigation of potential human host factors revealed mutations in the LARGE1 gene that may protect against severe Lassa fever in Nigeria (Kotliar et al., 2024). However, factors related to LASV evolution that could explain these differences have not been thoroughly investigated.

The Lassa virus genome is bi-segmented, coding for four proteins: the glycoprotein complex precursor, nucleoprotein (N), matrix (Z), and RNA-dependent RNA polymerase (L). The GPC is post-translationally cleaved into the Stable Signal Peptide (SSP), Glycoprotein subunit 1 (GP1) and Glycoprotein subunit 2 (GP2). Altogether, the GPC is crucial for cell entry, making it a key target for vaccine and therapeutic development (Garry, 2023; Katz et al., 2022). Despite its importance, our understanding of the LASV GPC, especially related to specific amino acid (AA) differences between lineages, is limited.

Knowing which lineage an emerging LASV outbreak strain belongs to provides immediate insight into the geographical origin of the outbreak, as distinct lineages are confined to specific locations. Moreover, since the lineages are known to elicit varying immunological responses (Buck et al., 2022) and exhibit different pathophysiological effects (Andersen et al., 2015), rapid lineage identification from a patient’s sample at the point of care would be highly advantageous. With the World Health Organization (WHO) supporting a significant expansion of genomic sequencing capabilities across Africa (Akande et al., 2023), genomic pathogen sequencing has the potential to become a critical component of outbreak response strategies. However, bioinformatic tools specifically designed for African pathogens, particularly the Lassa virus, are severely lacking.

In this study, we harness robust data, statistics, phylogenetics, and machine learning techniques to determine clade-defining amino acid mutations of possible clinical and evolutionary importance on the LASV GPC. We investigate the nature of important mutations and possible public health implications. Additionally, we introduce a rapid and accurate lineage assignment tool, which may enable healthcare professionals and researchers to quickly identify LASV lineages from sequence data. This tool may facilitate timely, targeted responses to both national and international Lassa fever outbreaks, improving surveillance, containment, and treatment strategies.

## Results

### Random forest classification reveals key LASV glycoprotein mutations

To investigate amino acid differences within the GPC of different lineages, we downloaded all available LASV nucleotide sequences released before December 1, 2023, and extracted and aligned the GPC region (supplementary figure 1). After quality filtering (see Methods), the final dataset contained 753 sequences, including 542 sequences from Nigeria, 141 from Sierra Leone, 11 from Guinea, 1 from Germany, 3 from Togo, 24 from Liberia, 5 from Mali, 13 from Côte d’Ivoire, and 13 from Benin. We translated the nucleotide sequences into amino acid sequences and verified the accuracy of the translation. We then grouped the sequences into two categories: those from Nigeria and those from other countries - consistent with Andersen et al. (Andersen et al., 2015).

To verify the positions identified by the RF method, we employed two additional techniques, Manhattan Distance (MD) and Pearson correlation (Pcorr). In this context, the MD is the sum of the absolute differences between AA positions across the two groups. We counted the occurrences of amino acids and gaps per position across the groups, normalized by sample number due to data imbalance, and computed the MD between corresponding position vectors. We ranked the positions and selected the top 15 for further analysis.

Finally, we applied Pearson correlation to assess variant similarity across the groups. Highly correlated positions suggest conservation, while low correlation indicates dissimilarity. We ranked the positions from least to most correlated and selected the top 15 for further analysis.

The application of these three methods revealed that 11 of the top 15 positions were consistently identified across all three methods (supplementary figure 3). Sorted from highest to lowest by cumulative ranking of all three sets, these were AA positions 273, 60, 61, 28, 44, 76, 74, 31, 421, 324, and 482. These positions are important in differentiating Nigerian lineages from other lineages. Some of the positions were previously found to be variable and involved in immune response (Wang et al., 2024; Ibukun, 2020). Positions 74 and 76, for example, have been implicated in the functioning of antibodies 20.10C and 36.1F (Enriquez et al., 2022). This suggests that the virus may evade these antibodies by varying these positions. Notably, deletion of amino acid positions 60-75, which includes 3 of the 11 most variable positions identified in our tests, has been shown to disrupt the functioning of all known anti-GPC antibodies (Robinson et al., 2016).

### An indel at amino acid position 60 shortens 98% of Nigerian GP1 sequences to 199AAs

The Manhattan Distance analysis highlighted position 60 as the most divergent between the groups (Figure 1B), and this position was consistently ranked highly by all three methods. Normalized value counts revealed that 98% of sequences from Nigeria have a gap at position 60, compared to less than 8.1% of sequences from other regions. This indel is located in the GP1 region, in the second position after the SSP cleavage site (Figure 2B). Consequently, at least 98% of Nigerian GPC sequences are shortened by one codon (nucleotide positions 178-180) at the nucleotide level and by one amino acid at the protein level compared to most sequences from other regions. This results in the GP1 of Nigerian sequences being 199 amino acids long compared to most sequences from elsewhere.

**Figure 2:**
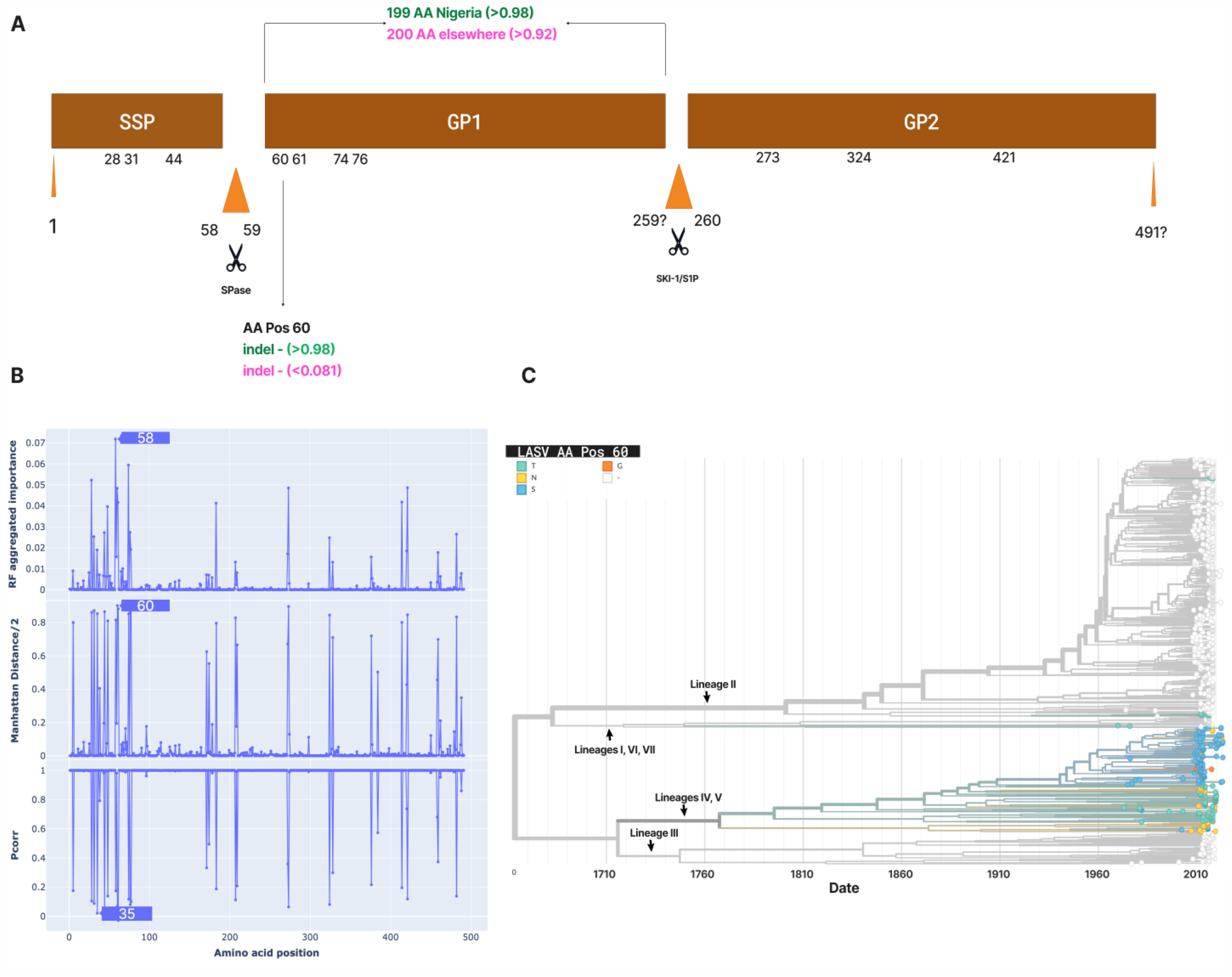
Differentiating positions on the LASV GPC (A) A representation of the most highly ranked positions jointly implicated by all three tests on the LASV GPC amino acid sequence. (B) Divergent regions identified by the random forest (top), Manhattan distance (middle), and Pearson correlation (Pcorr, bottom), respectively. (C) Time calibrated tree showing that the missing AA position 60 on Nigerian sequences is an insertion that predates the emergence of the Sierra Leonean strain. This insertion is seen to be present in all sequences in Lineage IV and Lineage V.

This discrepancy in length may affect protein folding and structure, potentially leading to functional and immunological variations. Additionally, the length inconsistency may cause issues with sequencing and assembly, as well as with polymerase chain reactions (PCR) targeted at the S segment or the GPC.

### Phylogenetic analysis suggests that position 60 constitutes an insertion that predates the emergence of Lineage IV, the Sierra Leonean strain

To investigate the nature of the indel at position 60 and its potential implications for public health, we reconstructed a time-scaled phylogenetic tree of the LASV GPC using the preprocessed data. The clock rate was estimated using Treetime (Sagulenko et al., 2018) at 8.20e-4 substitutions per site per year.

Phylogenetic analysis (Hadfield et al., 2018; Huddleston et al., 2021) revealed that the indel at position 60 is an insertion in Lineage IV, dating back to when this lineage first diverged from the Nigerian lineages (Figure 2C). The insertion is estimated to have occurred in the 18th century. All sequences on the major branch leading to lineages IV and V possess this insertion. The amino acids observed at this position include the Josiah genotype S60, with variants such as S60T, S60N and S60G. Only two sequences from Sierra Leone have the S60G variant (GenBank IDs OQ919514 and KM821773).

On the Nigerian side, lineages II and III almost entirely lack any amino acid at this position. This is consistent with the understanding that the Nigerian lineages are ancestral to the Sierra Leonean lineage (Andersen et al., 2015). Lineage VII, the Togo strain, which also includes sequences from the Benin Republic and a case imported to Germany in 2016, also lacks this insertion. However, a sequence closely related to Lineage VI (MK107927) has the insertion at position 60 (supplementary figure 5). Lineage I also have the insertion (a T60), as well as an amino acid deletion at position 62. Interestingly, differences in immune response have been reported for lineage I (Buck et al., 2022).

The AA 60 insertion appears to have recently reemerged in lineage II, particularly in sublineage IIb (Ehichioya et al., 2019), which is reported to circulate in the southern part of Nigeria (Figure 3). The affected states in Nigeria share close geographical boundaries. Meanwhile, the same insertion is also present in samples MH887782 and MH053506, which cluster closely with sublineage 2g, a sublineage with growing incidence in Edo and Ondo States in Nigeria (Happi et al., 2023) (supplementary figure 6). The temporal origin of these insertions was dated to around the last century.

**Figure 3:**
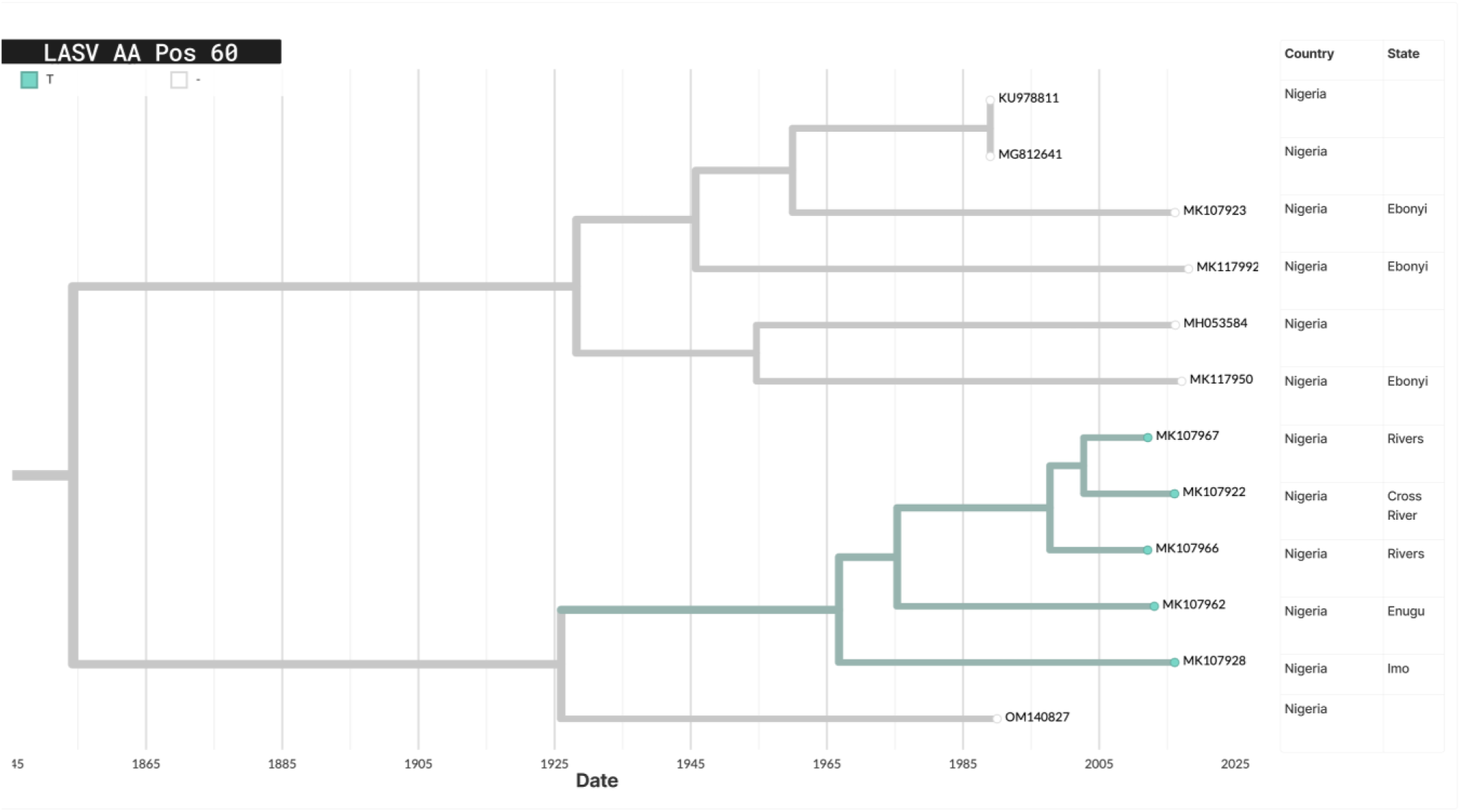
Re-emergence of the insertion on amino acid position 60 in lineage II. The most recent common ancestor (MRCA) that possessed this insertion emerged in the last century in the southern part of Nigeria.

The persistence of the insertion at AA position 60 over the years and across multiple lineages suggests that it may offer a fitness advantage for the virus. This could have significant implications for both the structural and functional aspects of the glycoprotein, potentially influencing virus transmission and virulence. The independent recurrence of insertions at this specific position further supports the idea that it provides a selective advantage to the virus. Additionally, it may point to possible adaptation to an intermediate host.

### Rapid LASV Lineage Classification from GPC sequence

Despite yearly Lassa fever epidemics, specific tools for its surveillance are scarce. Given that different LASV lineages circulate in specific regions and possess distinct clinical properties, the ability to efficiently assign unknown sequences to LASV lineages offers immense clinical and public health benefits. Consequently, we developed “CLASV”; a machine learning-based pipeline for LASV lineage assignment.

Using our curated data and workflow, we trained a Random Forest (RF) model to classify sequences into four groups: lineages II, III, IV & V, and VII. The training set included 480 sequences for lineage II, 59 for lineage III, 194 for lineages IV and V combined, and 16 for lineage VII. An initial test set, comprising 20% of each group and preprocessed in the same way as the training set, showed perfect performance across all evaluation metrics, with Accuracy, Precision, Recall, and F1-score all reaching 100%.

The model was subsequently exported into a pipeline and validated using newly published datasets from Adesina et al. (2023) and Bangura et al. (2024), which included 22 and 85 GPC sequences, respectively, as identified by our pipeline. These validation datasets were not used in the original training or testing phases, and the sequences from Adesina et al. (2023) were incomplete in terms of full GPC coverage. Nevertheless, CLASV (Figure 4) accurately classified sequences from both studies with 100% precision (Figure 5).

**Figure 4:**
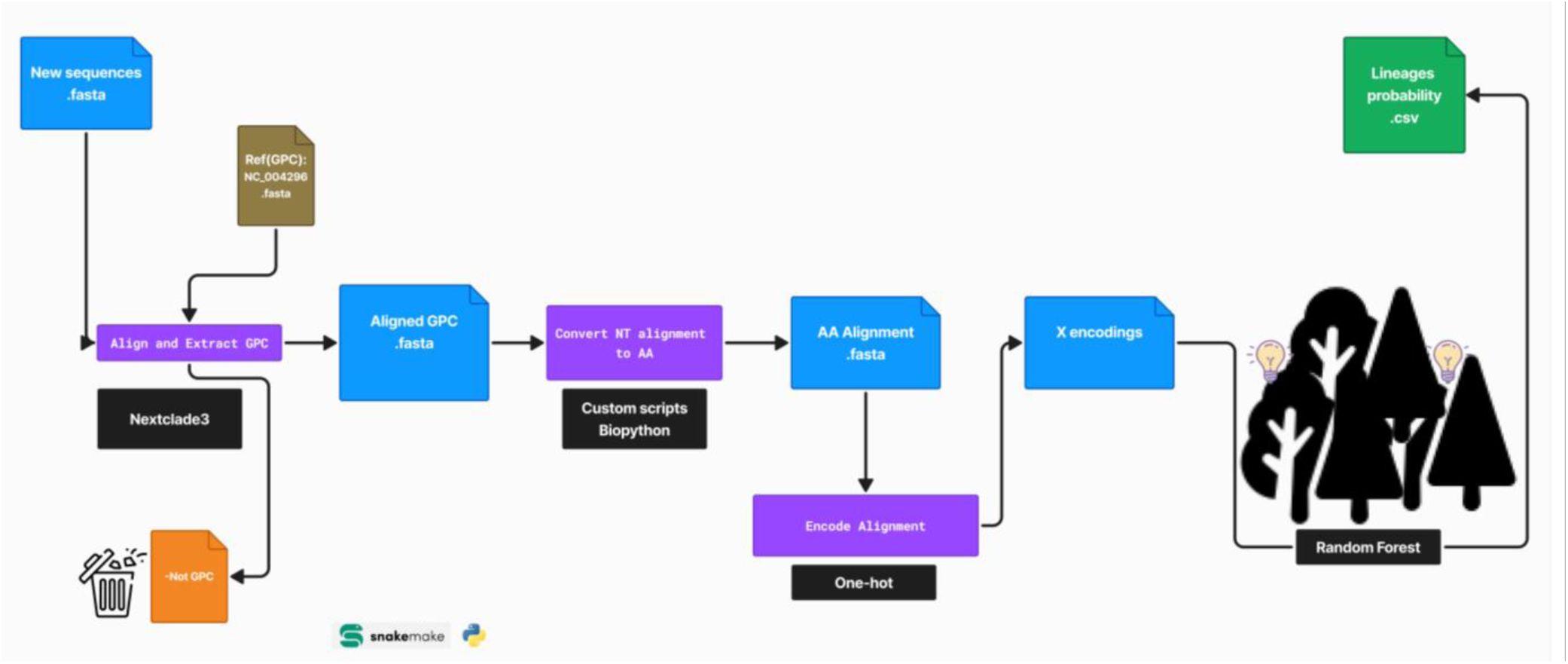
CLASV, a machine learning based pipeline for rapid LASV classification. The pipeline relies on the Snakemake engine for workflow management and Nextclade3, a part of Nextstrain (see Methods), as the GPC extractor and aligner. The aligned nucleotide sequences are converted into amino acid sequences and encoded similarly to the model’s training process. Subsequently, the encoded sequences are run through the trained random forest model, which predicts the probability per lineage. The highest probability is taken as the final prediction, provided it is greater than 0.5. Sequences with a probability below or equal to 0.5 are classified as inconclusive.

**Figure 5:**
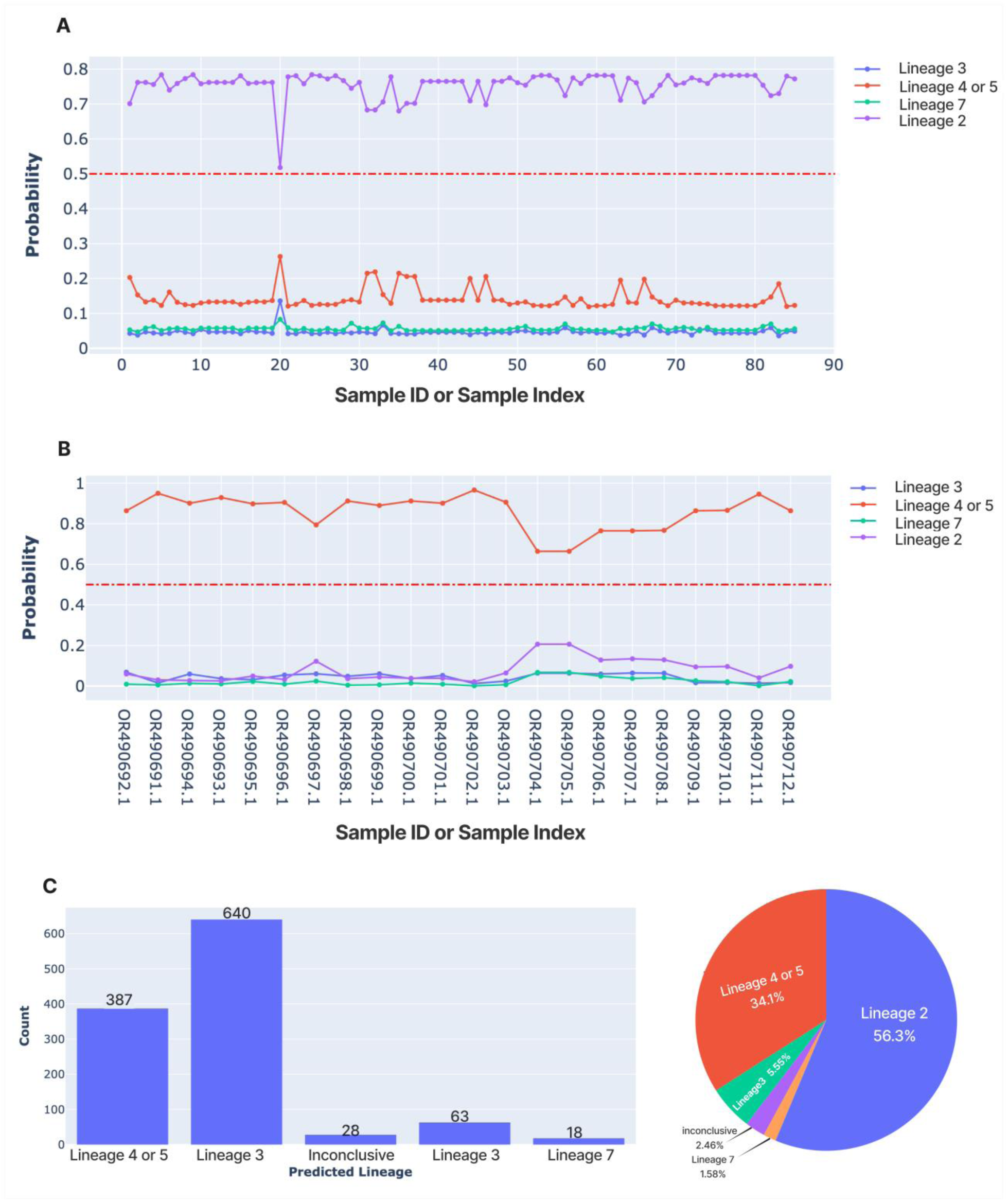
Performance of CLASV (A) CLASV accurately and precisely predicts sequences provided by Bangura et al. as belonging to Lineage 4 or 5. (B) The pipeline accurately and precisely predicts sequences provided by Adesina et al. as belonging to Lineage 2, despite the sequences being around half the size of the GPC. (C) The predicted distribution of LASV lineages in GenBank as of 06/07/2024. The inconclusive predictions include KM822128 (Pinneo), which belongs to lineage I. This reinforces the belief that lineage I is divergent from other lineages and emphasizes the pipeline’s ability to avoid false positives (supplementary GitHub code). Over half of all LASV GPC sequence in Genbank is predicted to belong to Lineage 2. These figures are auto-generated for every run, enabling rapid evaluation of the resulting lineage classification.

To test the pipeline’s speed, we downloaded all publicly available LASV sequences and processed them through the pipeline. On an average computer using five cores, it took less than two minutes to process and predict the lineages of all LASV GPC sequences in the dataset (Figure 5, supplementary GitHub code).

## Discussion

Lassa virus causes one of the deadliest infectious diseases, resulting in thousands of deaths annually. The virus comprises several lineages that circulate in different regions, with some exhibiting distinct immunological and virulent characteristics. In this study, we investigated the glycoprotein, which is the surface protein of LASV and the main target for vaccine and immunotherapy development. This protein is also important for drug development, and its nucleotide region is a key target for molecular diagnostics. We identified amino acid positions that are most variable by geographical location and lineage and investigated their prevalence. We also identified a recurring codon insertion that may confer a natural selection advantage to LASV. Finally, we provided a rapid, accurate, and easy-to-use lineage assignment tool for LASV sequences.

Andersen et al. (Andersen et al., 2015) reported differing pathophysiology and host codon usage between LASV lineages circulating in different regions. We have extended this work by identifying differing positions of the LASV GPC at the amino acid level. Some of the most divergent positions are around the cleavage site of the stable signal peptide by the signal peptidase, a protein reported to control the maturation of the GPC (Bederka et al., 2014). The variation in this region may lead to differences in viral replication and maturation.

Although the presence of a gap at amino acid position 60 in the LASV GPC has previously been reported (Buck et al., 2022; Perrett et al., 2023), its widespread prevalence had neither been clearly recognized nor demonstrated. We show that this gap is present in over 98% of Nigerian LASV sequences, including all lineage III sequences and nearly all lineage II sequences, making Nigerian lineages shorter by one amino acid compared to Sierra Leonean sequences (Lineage IV). To the best of our knowledge, this is the first time this has been categorically demonstrated. This prevalent variation could have profound implications for understanding the functional and phenotypic disparities between LASV lineages. Since the Josiah strain (Lineage IV), which is commonly used as a reference in the literature, contains 491 amino acids in the GPC, it is critical to highlight that the GPC of Nigerian lineages predominantly consists of 490 amino acids. This information is particularly relevant for protein engineering and LASV drug development, as the difference in length likely results in a difference in protein molecular weight. Such variations may affect protein structure, stability, and interactions, which are crucial for therapeutic and vaccine design.

Meanwhile, due to the divergent nature of LASV lineages, accurately aligning sequences to study genetic differences and virus evolution is challenging and often requires manual curation of alignments. For example, Wang et al. (2024) have aligned sequences as amino acid sequences before converting the alignment back to nucleotide sequences, presumably to avoid misalignment caused by sequence divergence. In a recent study Carr et al. (2024) reported deep mutational scanning of the Lassa virus GPC and published a phylogenetic tree in support of their results (https://nextstrain.org/groups/dms-phenotype/lassa/lassa-GPC?c=gt-Glycoprotein_60). Our results suggest that this tree may have been inferred from an incorrect alignment because in LASV strains that lack an amino acid at position 60, the authors filled the gap. This problem with studying indels may reflect a broader challenge in evolutionary biology, as highlighted by Redelings et al. (2024). In this study, we have provided and verified a LASV GPC alignment and showed evidence of important population-scale variations that should be considered in future research and development efforts for diagnostics, vaccine and immunotherapeutic development.

The effects of mutations are influenced by several factors, including the mutation’s position, the function of the affected protein, and the type of mutation. In multicellular eukaryotes, many traits are polygenic, so a single mutation may have little to no observable effect. However, individual mutations can have severe consequences for monogenic traits. For instance, in sickle cell disease, a single nucleotide substitution results in an amino acid change, which can have catastrophic effects when both gene copies are affected (Inusa et al., 2019). In viruses, which have smaller genomes and fewer proteins, mutations can also have profound effects, particularly when they occur in critical proteins. This is especially true when the affected protein is the virus’s sole surface protein, as is the case with LASV GPC. Indeed, a well-known example of the impact of this kind of viral mutation is the D614G substitution in the spike glycoprotein of SARS-CoV-2, which was reported to confer a selective advantage by increasing the virus’s binding affinity to the angiotensin-converting enzyme 2 (ACE2) receptor, making it the dominant variant at the time (Rao et al., 2021). Similarly, indels in the spike glycoprotein of SARS-CoV-2 were implicated as mechanisms of selective advantage (Zhou et al., 2021). The insertion at amino acid LASV GPC position 60 could, therefore, be a major contributor to differences in pathophysiology and immune response among LASV lineages. This mutation occurs in GP1, the subunit responsible for binding the LAMP1 and alpha-dystroglycan receptors during viral entry (Cao et al., 1998; Jae et al., 2014), potentially altering cell entry efficiency between Nigerian lineages and those from other regions, such as the Sierra Leonean strain. Interestingly, Andersen et al. (2015) observed that the genome abundance of the Sierra Leonean strain in patients is significantly higher than that of Nigerian lineages. Since viruses depend on host cellular mechanisms for replication, enhanced cell entry—possibly influenced by the insertion in GP1—might contribute to this higher genome abundance. The viral load increase, suggested by the higher genome abundance, could explain the greater fatality rates associated with the Sierra Leonean strain compared to its Nigerian counterparts.

We demonstrated that the indel at amino acid position 60 is an insertion that predates the divergence of the Sierra Leonean lineages from the Nigerian lineages. All sequences from lineages IV and V in our dataset possess this insertion. Additionally, there has been an independent re-emergence of this insertion at the same position in lineage II in recent years. The persistence of this mutation strongly suggests that it provides a fitness advantage through natural selection. Indeed, the findings of Andersen et al. (2015) indicate that the Sierra Leonean strain, which predominantly harbors this insertion, exhibits better survival in human hosts. From a public health perspective, monitoring this insertion is crucial, as it represents a relatively unique mutation in the GPC, and its recurrence in lineage II raises concerns. Hypothetically, the possibility that an unknown intermediate host may be facilitating this insertion cannot be ruled out. If the virus enters such a host, this mutation could play a significant role in viral replication and adaptation. However, it is clear that the insertion has emerged, re-emerged independently, persisted, and reached the population level, indicating its potential viability. A key area for future research concerns the exact functional implications of this recurring insertion. This will deepen our understanding of inter-lineage differences and the evolutionary trajectory of LASV, which are critical for informing surveillance efforts, vaccine development, and immunotherapy strategies.

Given the large prevalence of the codon insertion at position 178-180 of the GPC, it is advisable to avoid targeting this region with PCR primers, as this could result in inconsistent results across different LASV lineages due to variability in binding affinity depending on the reference sequence used for primer design. Similarly, the design of ELISA (Enzyme Linked Immunosorbent Assay) test kits should take into account the variability at this amino acid position (LASV GPC 60) to avoid inaccuracies in detecting the virus. The presence of this insertion can affect the epitope structure, potentially leading to reduced recognition by antibodies used in ELISA, which are designed based on a reference sequence that may not account for such variations. Interestingly, there has been evidence of PCR false negatives for Lassa virus which were detected by metagenomics sequencing (Kotliar et al., 2024). PCR false negatives are especially critical when dealing with a BSL-4 pathogen with pandemic potential such as LASV (WHO, n.d.). A resulting incorrect diagnosis could result in a patient’s death, further local transmission, and even the introduction of the pathogen to other continents. Therefore, it is essential to design diagnostic test kits that are either robust to inter-lineage differences or specific to the respective lineages to ensure accurate and reliable results.

Our finding that positions 60, 61, and 74 are among the top positions varying between lineages is consistent with their established role in immune evasion (Robinson et al., 2016). Supported by reports from Enriquez et al. (Enriquez et al., 2022) and Carr et al. (Carr et al., 2024), monoclonal antibody 21.10C may be ineffective in Southern Nigeria due to the high prevalence of the immune-resistant mutation at position 76 (supplementary figure 7). Monitoring these positions is crucial, alongside others such as position 273, which has also been identified as immunologically significant (Ibukun, 2020; Robinson et al., 2016) and consistently ranked as highly variable in our analyses.

In addition to LASV, Africa hosts several other endemic pathogens, such as Ebola virus, Marburg virus, and Crimean-Congo hemorrhagic fever virus. While the WHO made significant strides in enhancing surveillance capacity in Africa during the COVID-19 pandemic (Akande et al., 2023), there remains a critical need for continuous improvements. One of the major areas requiring further development is bioinformatics capacity, which remains insufficient across much of the continent (Sharaf et al., 2023). Given the recurring LASV epidemics (Garry, 2023), there is an urgent need for specialized tools that facilitate rapid responses. To address this, we developed a pipeline for rapid LASV lineage assignment, employing a method similar to that used in our motif search (Figure 6).

**Figure 6:**
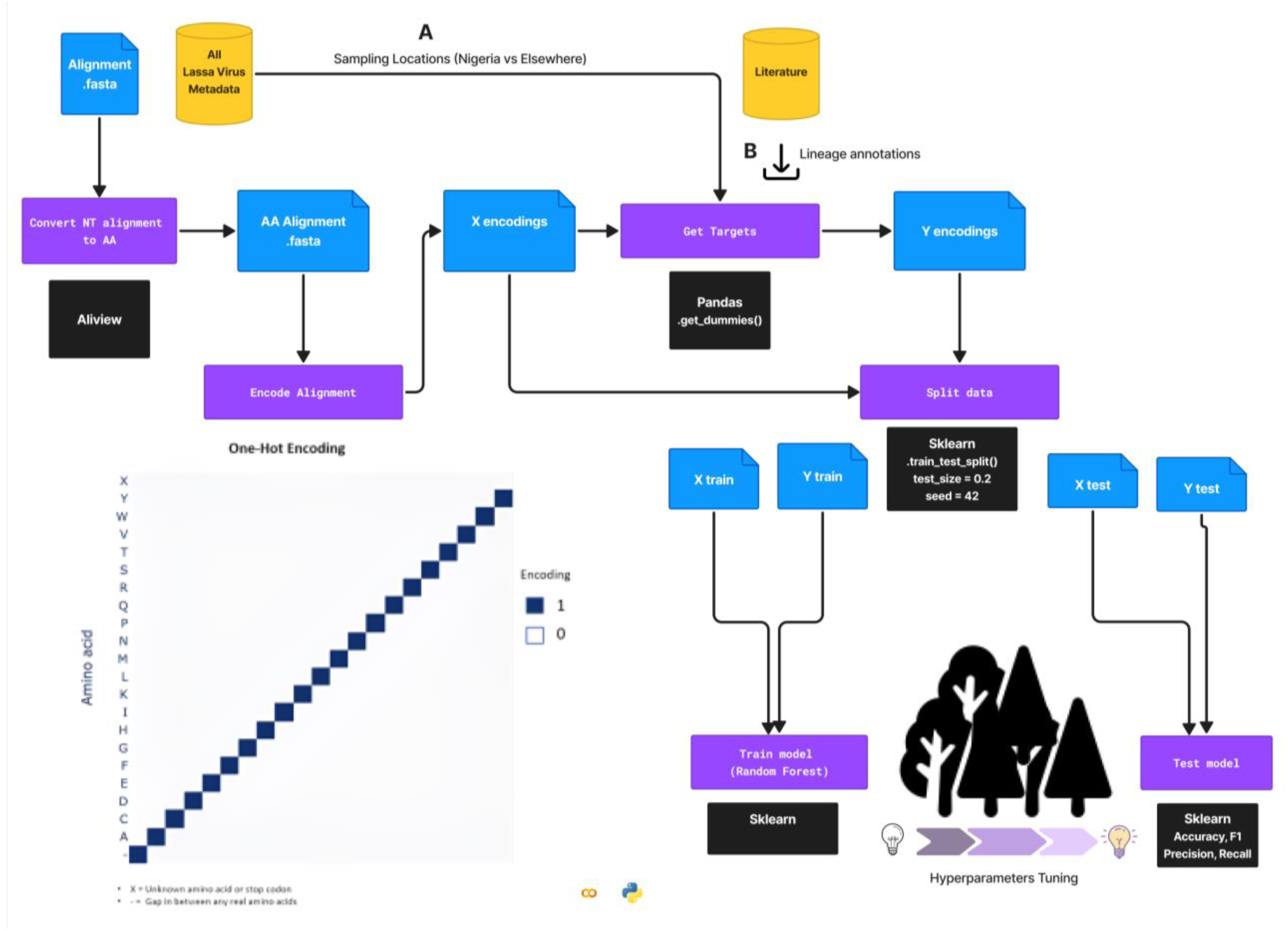
Motif finding and classification workflow An overview of the machine learning workflow demonstrates how the two Random Forest models were created using a similar process. (A) The sampling locations were annotated using metadata. (B) The lineages of each sequence in our dataset were annotated using insights from the literature. To achieve optimal performance from the models, hyperparameter tuning, primarily involving varying the number of decision trees, was conducted.

Our machine learning-based pipeline, CLASV, offers a novel capability for researchers and clinicians to rapidly assign lineages to LASV sequences. With built-in data cleaning processes, high speed, precision, and ease of use, this tool is invaluable for managing endemic Lassa fever cases in West Africa. Additionally, the pipeline can assist in managing imported cases and contribute to global public health efforts. The ability to quickly assign lineages offers immediate insights into the origin of a Lassa fever case, informs treatment decisions, and facilitates a more effective response. Developed in Python programming language and based on Snakemake (Köster & Rahmann, 2012), CLASV can be easily integrated into other workflows, enhancing its utility in both research and clinical settings. Machine learning methods are generally faster than traditional approaches, especially in genomic sequence lineage assignments, where speed is crucial during an outbreak.

A limitation of the pipeline is the necessity for regular updates as new lineages emerge. Currently, CLASV can reliably assign the two major lineages, lineage II and lineage IV, as well as lineages VII and III. However, as more data becomes available, the model can be seamlessly updated using the workflow we have established. This tool represents a significant contribution a widely recognized need, the strengthening of bioinformatics capacity on the African continent (Nembaware et al., 2023; Olono et al., 2024; Sharaf et al., 2023).

Finally, we highlight the need for more sequencing to properly study LASV evolution and to develop additional tools. The establishment of standard guidelines for new LASV lineage naming, as suggested by (Whitmer et al., 2018), would help make tools like this pipeline as reproducible and reusable as possible.

## Methods

### General preprocessing

LASV sequences released between December 1905 and December 1, 2023, along with their accompanying metadata, were downloaded from NCBI Virus (available at https://www.ncbi.nlm.nih.gov/labs/virus/vssi/#/). To ensure the inclusion of only field samples, the ’exclude lab strain’ filter was applied. Using the GPC gene from the reference ID NC_004296, we extracted and aligned the GPC regions from all sequences using LAST (Kiełbasa et al., 2011) and MAFFT (Katoh et al., 2002), available at https://mafft.cbrc.jp/alignment/server/specificregion-last.html, resulting in 1021 GPC sequences. Sequences with more than 5% gaps and ambiguous nucleotides of the total alignment length were removed, reducing the dataset to 808 sequences. The final stop codon position was removed from the alignment because the stop codon signals the termination of translation, and it does not encode an amino acid. Sequences lacking sampling dates and locations were excluded, leaving 753 sequences.

Alignment visualization and exploration were conducted using Aliview (Larsson, 2014). Manual curation was performed to ensure codon consistency. Specifically, misalignment involving codons disrupted by 3 gaps was adjusted by moving a single nucleotide to match the two others to ensure proper translation. Nucleotide to amino acid translation was done using Aliview.

All data analyses were conducted in Python using standard libraries such as pandas (McKinney, 2010) (available at https://pandas.pydata.org/), scikit-learn (Pedregosa et al., 2011), and Biopython (Cock et al., 2009). The workflow was implemented using Jupyter Notebook in Google Colab (Bisong, 2019).

### Phylogenetic analysis of GPC

Using a pipeline based on Snakemake (Köster & Rahmann, 2012), Nextstrain (Hadfield et al., 2018), and Augur (Huddleston et al., 2021), we reconstructed both a Maximum Likelihood tree and a time tree (supplementary Figure 9). The alignment and metadata, which includes sampling dates, country, and host information, are the inputs to the pipeline. A Maximum Likelihood tree was built using IQ-TREE (Nguyen et al., 2015) with the application programming interface (API) provided by Augur. Similarly using Augur API, the tree was processed by TreeTime (Sagulenko et al., 2018), along with the metadata, to generate a time-calibrated tree.

The trees and accompanying metadata were parsed using the Augur export command into a JSON file, which was subsequently visualized using Auspice (available at https://auspice.us/), a part of the Nextstrain (Hadfield et al., 2018) toolkit. All phylogenetic images were generated using Auspice - which have been edited for clarity in Figma (available at https://www.figma.com/).

### Sequence annotation

Sampling locations were extracted from the accompanying metadata for each sequence from GenBank. Sequences were annotated based on the country column to analyze motif differences between Nigerian sequences and those from other countries. The sequences were then grouped into those from Nigeria (542 sequences) and those from other countries (211 sequences).

For lineage annotation, clades in our reconstructed phylogenetic tree were annotated based on information from the literature (Andersen et al., 2015; Ehichioya et al., 2019; Manning et al., 2015; Olayemi et al., 2016; Whitmer et al., 2018; Yadouleton et al., 2020) (see supplementary GitHub data). The leaves of each clade were then collected. Specifically, the sequences were categorized into the following lineages: 480 sequences for lineage II, 59 sequences for lineage III, 194 sequences for lineages IV and V combined, and 16 sequences for lineage VII. Lineages I and VI were excluded due to insufficient data. Lineages IV and V were grouped together based on the recommendation of Whitmer et al. (Whitmer et al., 2018), who pointed out that the distance between the two lineages is similar to those between other sublineages.

### Machine learning: Random forest

#### Sequence encoding

The amino acid alignment, comprising 753 sequences and 491 positions, was loaded using the Biopython alignment class. Gaps preceding and following any real amino acids were converted to ‘unknowns’, as these gaps typically indicate either short sequences or sequencing issues and therefore lack biological significance.

To prevent any biases, all features and targets were one-hot encoded. Each base of every amino acid sequence in the alignment was encoded into a 21-position vector. Gaps between bases were similarly encoded, while ‘unknowns’ were encoded with zero values. The encoded features were then flattened, with each position in the vector transformed into an individual column, resulting in a data matrix of dimensions 753 x 10,311.

The targets were encoded using one-hot variables, facilitated by pandas’ `get_dummies()` method. We trained two models with different targets: country and lineage.

#### Model training

The dataset was split into training and testing sets using scikit-learn with a random state of 42, ensuring that the test set comprised 20% of the total dataset and stratification was based on the target variable. Using the training set, we trained Random Forest (RF) models with the scikit-learn package. The RF model was chosen for its robustness in handling multidimensional and imbalanced data. The random state for the RF model was set to 80, and the seed state was set to 42 to ensure reproducibility. A total of 100 decision trees were used in the binary country classification, while a total of 1,000 decision trees were used in the lineage classification.

#### Model Evaluation

The models’ performance was evaluated using the test dataset, which comprised 20% of the entire dataset. Standard metrics from the scikit-learn package, such as precision, F-score, and recall, were used to assess the models’ performance through the `classification_report()` function.

Since amino acid sequences can be identical between samples due to extreme conservation, we also investigated possible overlapping data points between the training and test sets. Approximately twenty percent of the test data was found to be present in the training set. However, this overlap does not affect the results, as the model achieved 100% across all test metrics.

#### Feature importance selection

During the encoding process, each position was represented by a 21-vector, and the resulting matrix was flattened, creating 491 x 21 columns (totaling 10,311 columns). Therefore, each position in the amino acid sequence is represented by 21 features.

The Random Forest algorithm outputs a large matrix containing the column index and the feature importance (10,311 rows by 2 columns). To determine the exact position of an amino acid in the alignment, each column index is divided by 21, and the integer part of the quotient is taken and incremented by 1. The weights of each position were combined into an aggregated feature importance score. Each feature corresponds to an exact position in the GPC amino acid sequence. The positions were subsequently ranked using the aggregated feature importance score from highest to lowest.

### Manhattan Distance and Pearson Correlation

#### Manhattan Distance

Given an amino acid alignment with sequences belonging to either of two categorical variables, it is possible to investigate positions of differences this way:

**Group the AA alignment into two categories based on metadata:** Nigeria (542 sequences) and Elsewhere (211 sequences).

**Calculate amino acid counts and gaps per position for both groups:** Resulting in 21 by 491 matrices for each group (which is the 20 amino acids + gap (21) by length of the alignment (491))

#### Preprocessing

● Convert gaps before and after any real amino acids to “Unknowns”.
● Add the “Unknowns” count per position to the most frequent variant in that position.

**Normalize each matrix:** Divide by the number of samples.

**Compute Manhattan distance:** Calculate the absolute distance between the groups based on the normalized matrices as follows:

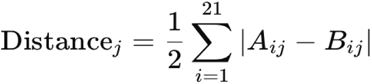

where A is the normalized amino acid count matrix of Nigerian sequences, B is the normalized amino acid count of sequences from other countries (Elsewhere), and 2 is the maximum absolute distance per position. The resulting vector contains the sum of the absolute differences for each position which was then ranked from highest to lowest (supplementary figure 8).

#### Pearson correlation

The normalized amino acid counts of the Nigeria and Elsewhere groups were correlated using the normalized matrices from the previous step. Using the pandas.corr() method, we calculated the Pearson correlation between each corresponding column of the Nigeria and Elsewhere matrices. This calculation produced a correlation vector representing each amino acid position.

The resulting vector, containing the correlation score for each position, was then ranked from least to highest by the absolute correlation score. This ranking highlights the positions with the most significant differences in variability between the two groups.

## Data and code availability

**Supplementary GitHub Data:** https://github.com/JoiRichi/LASV_ML_manuscript_data

**Supplementary GitHub code (Machine learning):** https://github.com/JoiRichi/CLASV

**Supplementary Figures:**

Attached

## Acknowledgement

This work is funded by the federal Government of Germany.

We especially and wholeheartedly thank the Nextstrain main developers (https://nextstrain.org/team) for providing open-source code for phylogenetic analysis and for supervising the modification for Lassa virus. We also thank John Huddleston, Joseph Prescott, Zewen Yang, Sodiq Ayobami Hameed, Manfred Weiss, Laila Benz, and Knut Reinert for their helpful comments.

