## Supplementary Figures for "Machine Learning Reveals Key Glycoprotein Mutations and Rapidly Assigns Lassa Virus Lineages" (from another version of this preprint: version 1, 2024-07-31)

Further information available at:

https://github.com/JoiRichi/LASV_ML_manuscript_data

Figure 1: Data preprocessing

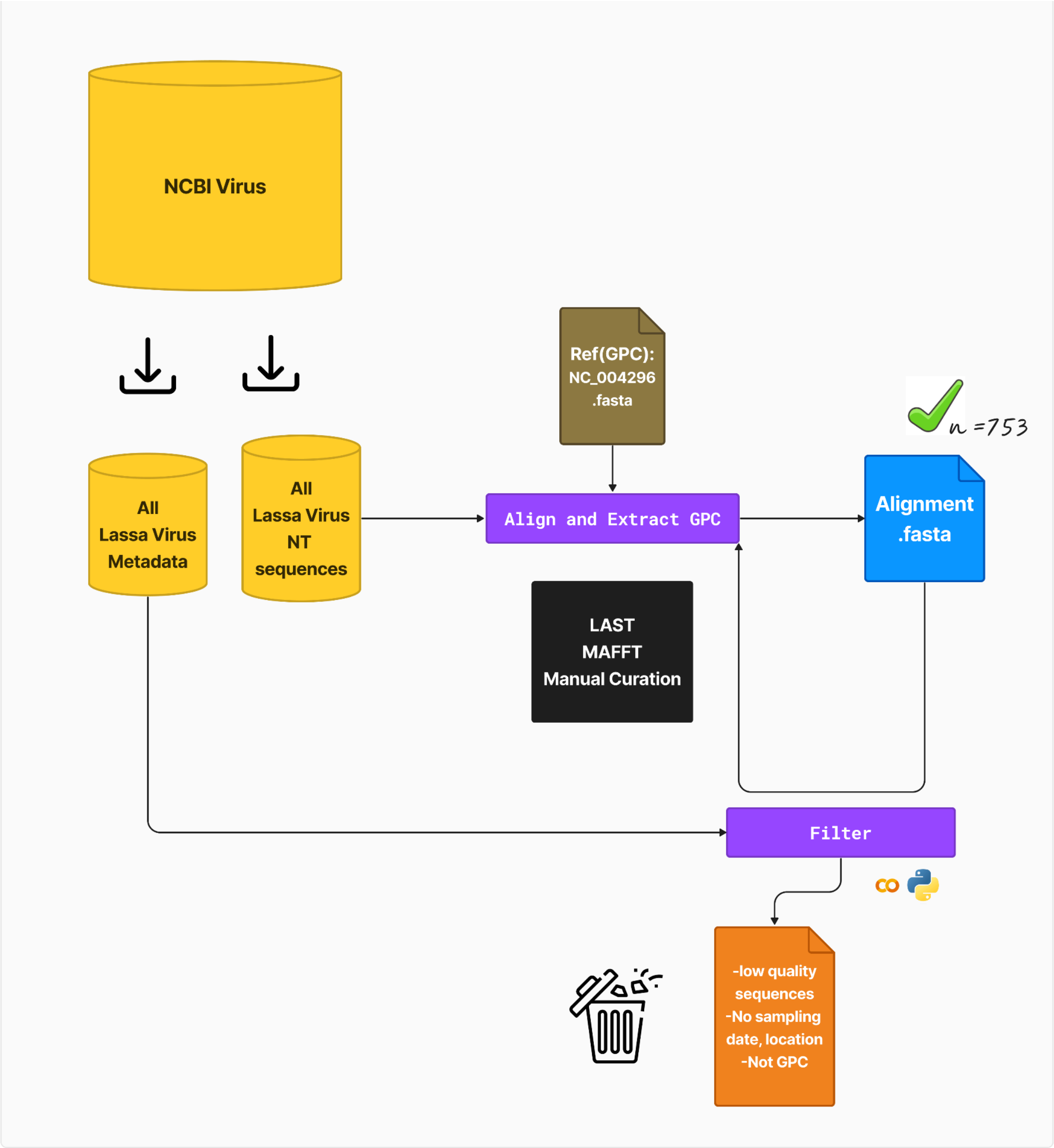


Figure showing the preprocessing pipeline. 753 sequences were proceeded to further analysis.

Figure 2: Histogram of random forest feature aggregate importance scores

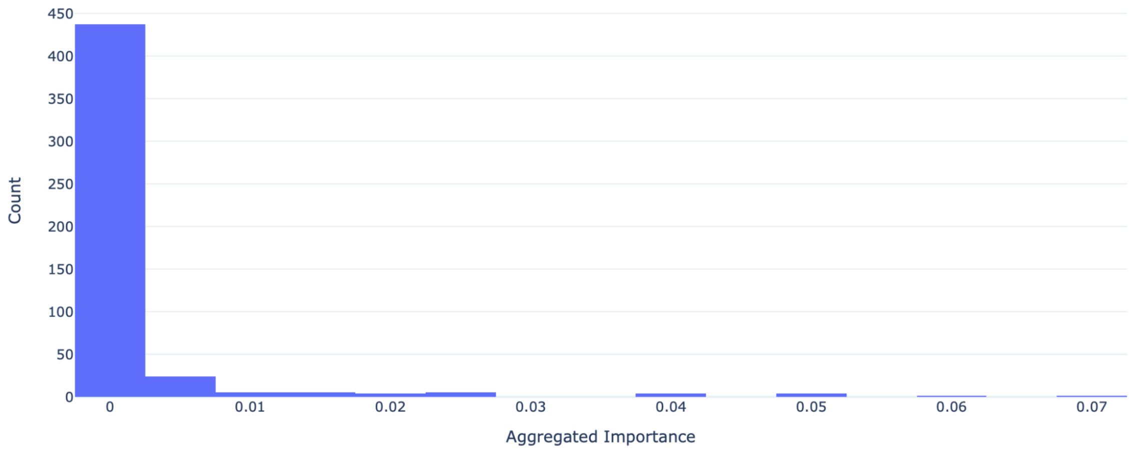


Very few positions are different between glycoprotein of Lassa virus (LASV) circulating in Nigeria and those from other countries, as implicated by the random forest classifier.

Figure 3: Most variable positions on the LASV glycoprotein complex (GPC)

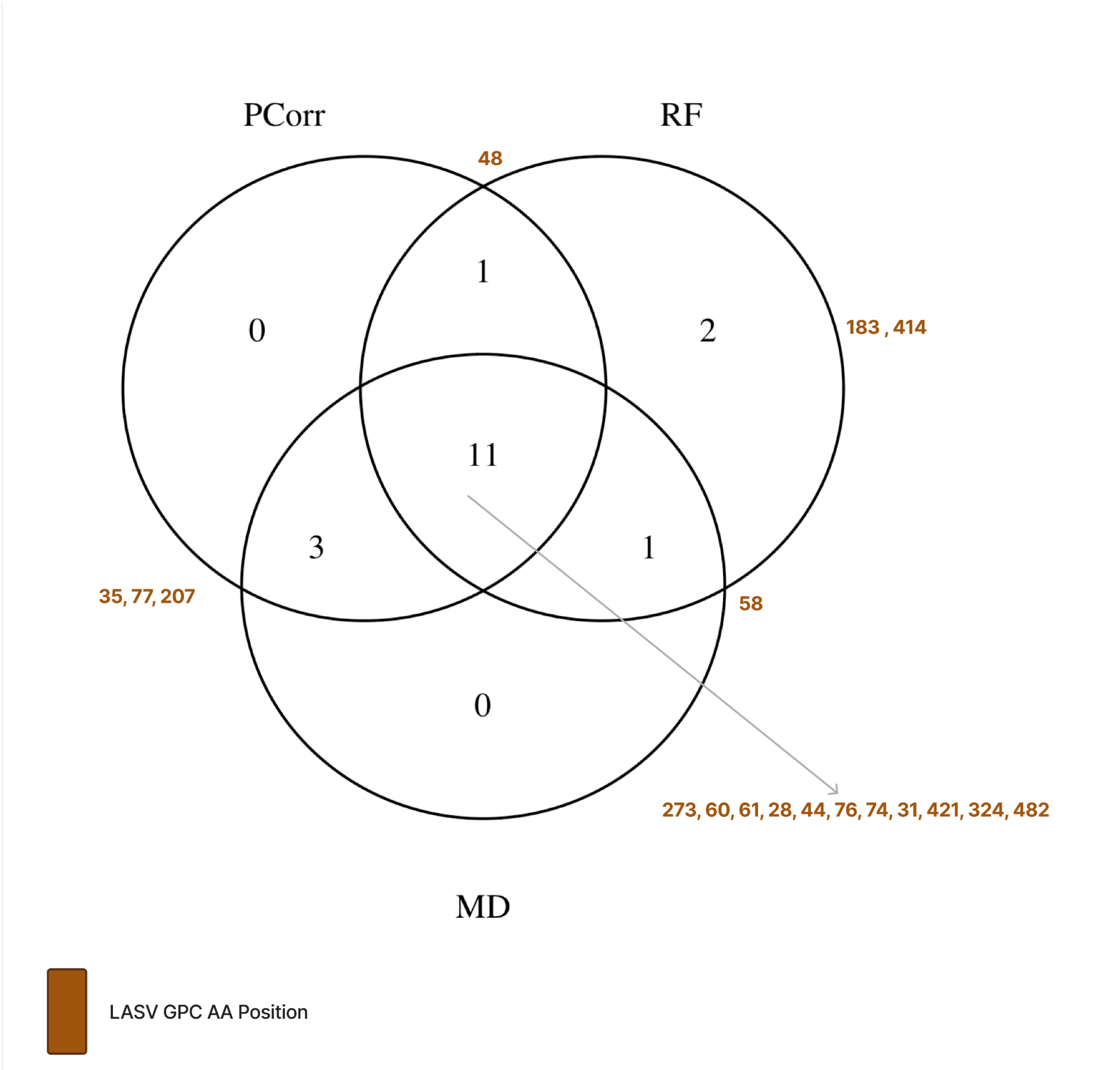


Venn diagram showing the overlap among the top 15 positions selected from the outputs of three tests: Random Forest (RF), Pearson Correlation (PCorr), and Manhattan Distance (MD). Eleven (11) of these positions were jointly implicated by all three methods. The 11 positions have been sorted using their cumulative rank across the three tests.

Figure 4: Distribution of amino acid at LASV GPC position 60

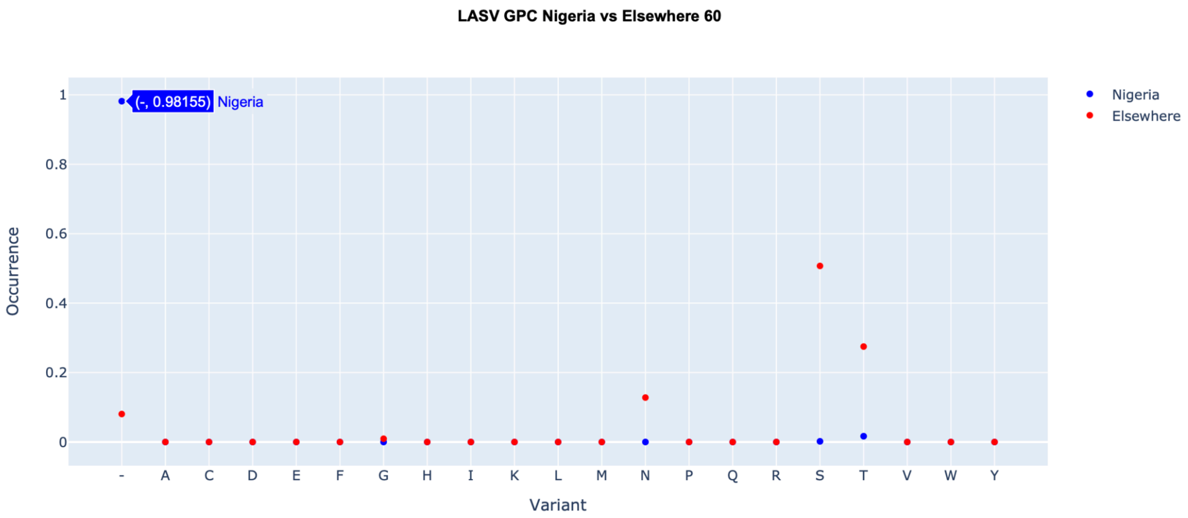


Normalized value count reveals that over 98% of Nigerian LASV GPC sequences have an indel on position 60.

Figure 5: Indel in lineages VII, VI and I at LASV GPC position 60

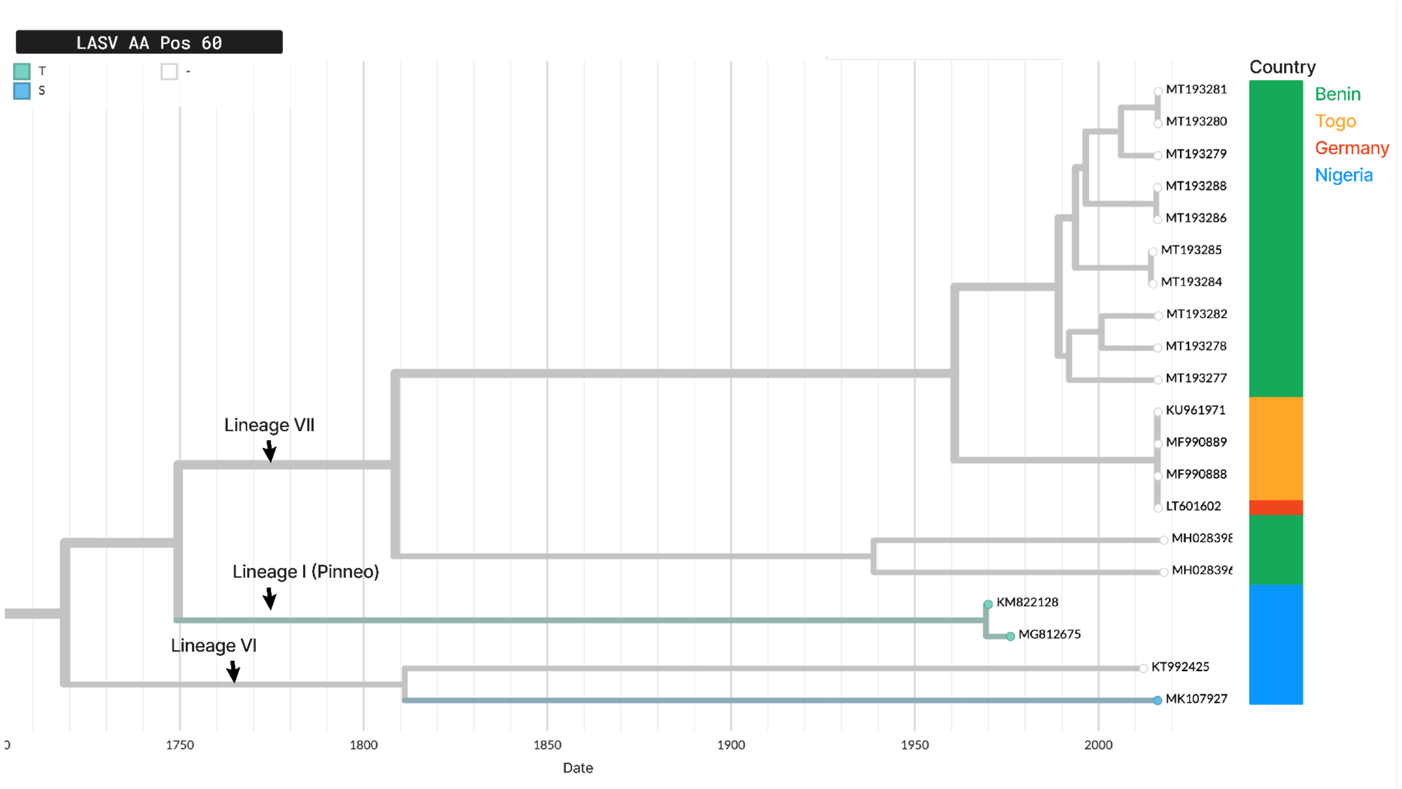


Lineage VII, the Togo strain, which also includes sequences from the Benin Republic and a 2016 imported case to Germany, lacks this insertion, similar to lineages II and III. However, a sequence closely related to Lineage VI (MK107927) is seen to have the insertion at position 60. Lineage I, on the other hand, does have the insertion, a T60.

The tree topology closely resembles previous results (Whitmer et al., 2018).

Figure 6: Re-emergence of insertion in sublineage IIg LASV GPC

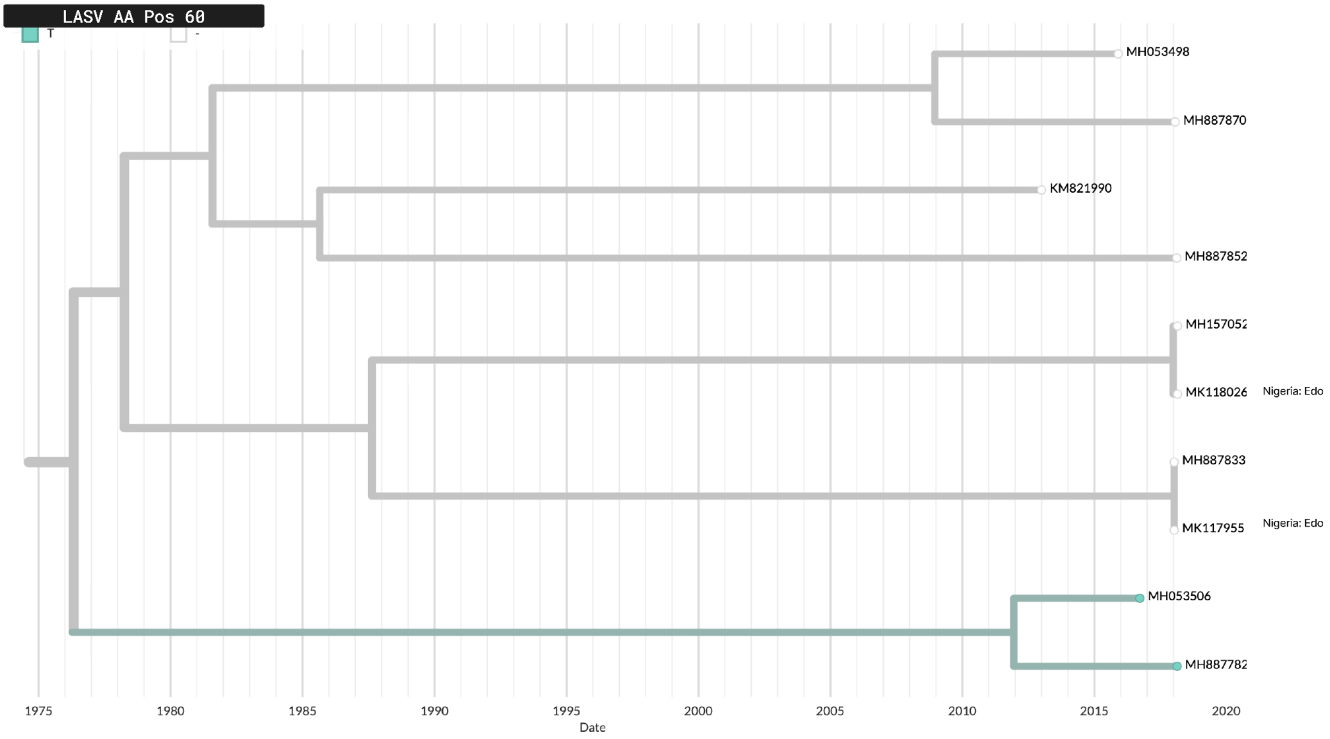


The insertion seems to have reappeared in sequences closely related to sublineage IIg at position 60, a sublineage with growing incidence in Edo and Ondo States in Nigeria (Ehichioya et al., 2019; Happi et al., 2023).

Figure 7: Massive prevalence of implicated immune resistant Mutation in lineage II LASV GPC

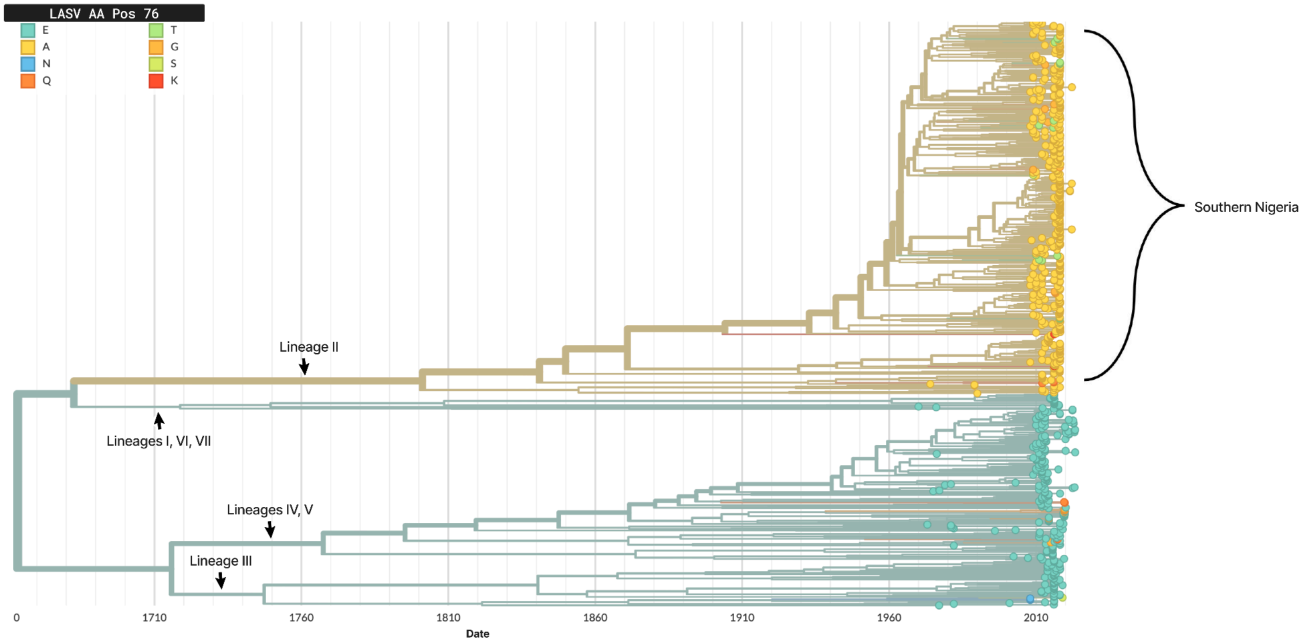


Figure showing the massive prevalence of amino acid A76 in lineage II. Lineage II is known to circulate in Southern Nigeria (Ehichioya et al., 2019). Supported by reports from Enriquez et al. (Enriquez et al., 2022), monoclonal antibody 21.10C may be ineffective in Southern Nigeria due to the high prevalence of the immune-resistant mutation at position 76.

Figure 8: Manhattan Distance and Pearson correlation

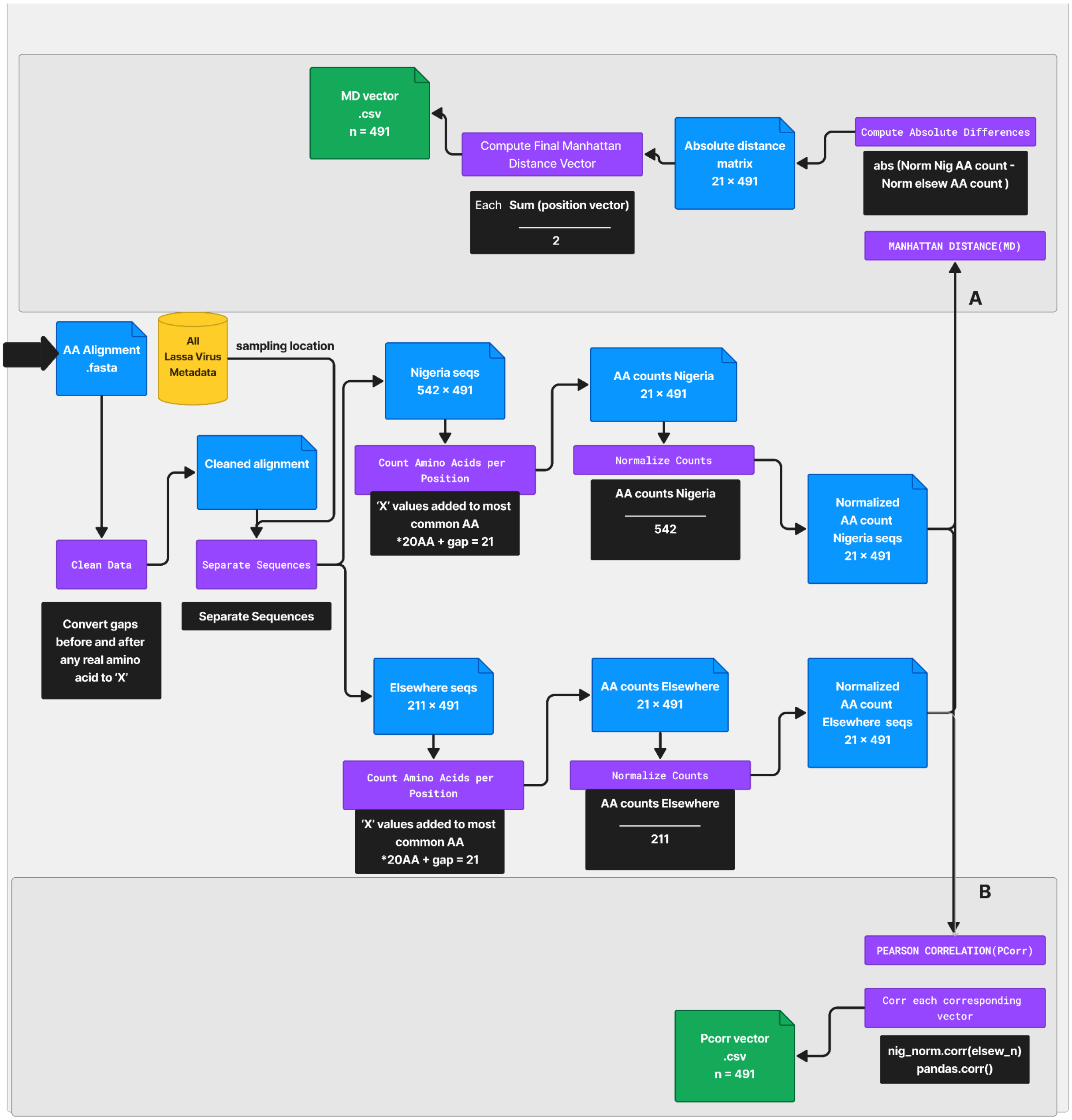


Figure showing the workflow of the Manhattan distance and Pearson correlation analyses. The preprocessed alignment serves as the input, and both methods share similar steps until the normalized matrices are obtained for the two groups (Nigeria vs. Elsewhere). Subsequently, the distance is calculated in step (A) while the correlation is calculated in step (B).

Figure 9: Figure showing the phylogenetics pipeline

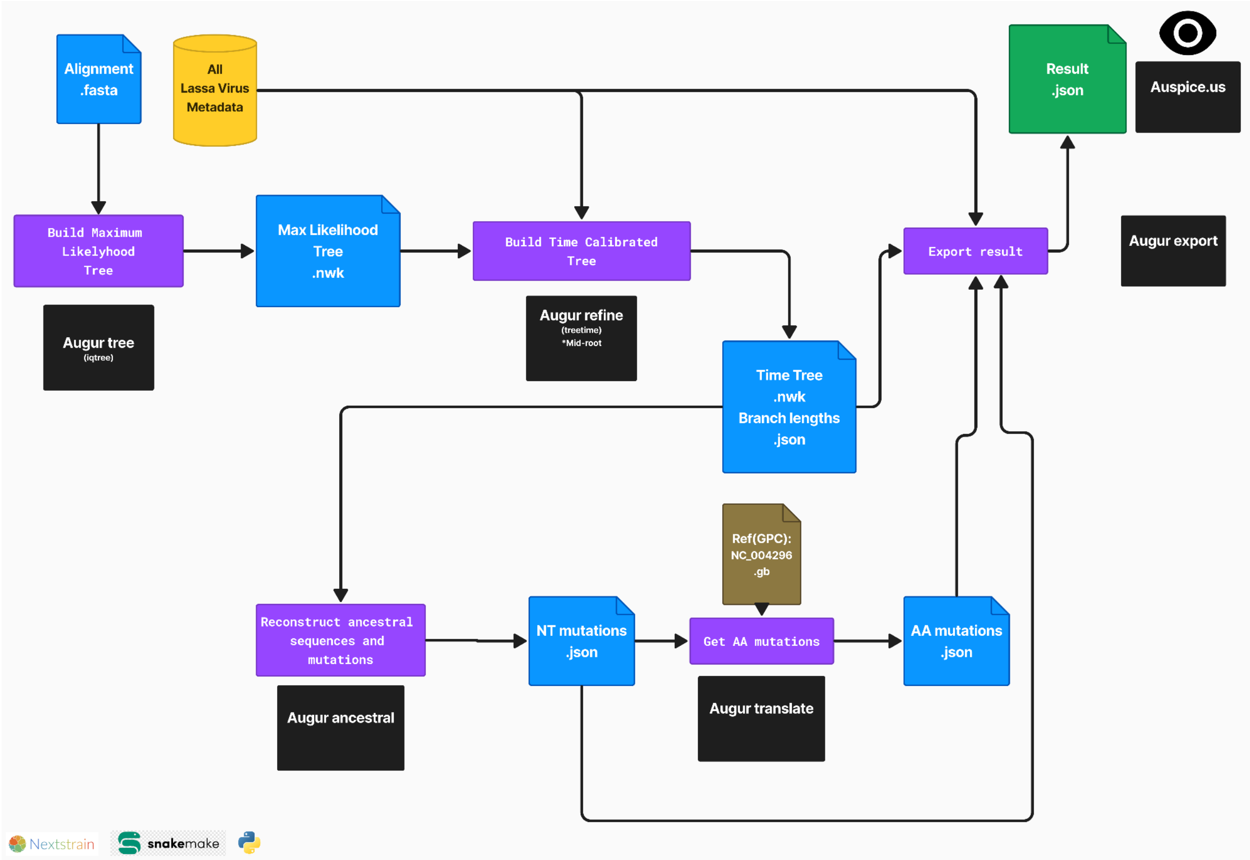


Using a pipeline based on Snakemake, Nextstrain, and Augur, we reconstructed both a Maximum Likelihood tree and a time tree. The pipeline starts with an alignment and metadata, including sampling dates, country, and host information. A Maximum Likelihood tree was constructed using IQ-TREE with the Augur API. Subsequently, the tree was processed by TreeTime, along with the metadata, to generate a time-calibrated tree. The trees and accompanying metadata were exported into a JSON file and visualized using Auspice.
